# A *Drosophila* Model for Behavioral Sleep Modification

**DOI:** 10.1101/391375

**Authors:** Samuel J. Belfer, Alexander G. Bashaw, Michael L. Perlis, Matthew S. Kayser

## Abstract

Insomnia is the most common sleep disorder among adults, especially affecting individuals of advanced age or with neurodegenerative disease. Insomnia is also a common comorbidity across psychiatric disorders. Cognitive Behavioral Therapy for Insomnia (CBT-I) is the first-line treatment for insomnia; a key component of this intervention is restriction of sleep opportunity, which optimizes matching of sleep ability and opportunity, leading to enhanced sleep drive. Despite the well-documented efficacy of CBT-I, little is known regarding how CBT-I works at a cellular and molecular level to improve sleep, due in large part to an absence of experimentally-tractable animals models of this intervention. Here, guided by human behavioral sleep therapies, we developed a *Drosophila* model for behavioral modification of sleep. We demonstrate that restriction of sleep opportunity through manipulation of environmental cues improves sleep efficiency and quality in multiple short-sleeping *Drosophila* mutants. The response to sleep opportunity restriction requires ongoing environmental inputs, but is independent of the molecular circadian clock. We apply this sleep opportunity restriction paradigm to aging and Alzheimer’s disease fly models, and find that sleep impairments in these models are reversible with sleep restriction, with associated improvement in reproductive fitness and extended lifespan. This work establishes a model to investigate the neurobiological basis of CBT-I, and provides a platform that can be exploited towards novel treatment targets for insomnia.

## Introduction

Insomnia is the most common sleep disorder among adults, with significant public health and economic consequences^1–4^. Cognitive Behavioral Therapy for Insomnia (CBT-I) is the first-line intervention for treatment of insomnia^5^. CBT-I includes a combination of modalities: behavioral therapy (restriction of sleep opportunity and stimulus control), cognitive therapy (cognitive restructuring of dysfunctional beliefs about sleep and sleep disturbances), and sleep hygiene (education pertaining to behaviors that facilitate sleep continuity). Recent work suggests that restriction of sleep opportunity alone is sufficient to gain most of the benefits of CBT-I^6^. Sleep restriction addresses a prominent clinical feature of insomnia: the mismatch between sleep opportunity and sleep ability. Patients with insomnia often expand time in bed (sleep opportunity extension) with the goal of recovering lost sleep^7^. This adaptation is thought to perpetuate insomnia in the long term by promoting the mismatch between sleep ability (low) and opportunity (high), leading to less efficient, less consolidated sleep. By restricting time in bed, sleep restriction optimizes matching of sleep ability and opportunity, leading to enhanced sleep drive (increased homeostatic pressure for sleep) and more consolidated sleep. Sleep opportunity is titrated as sleep ability stabilizes and increases. Although CBT-I has shown reliable and durable efficacy for insomnia treatment^8–10^, limited accessibility of practitioners and long duration of therapy are obstacles to broad implementation^11–13^. If behavioral sleep interventions could be studied at a molecular/cellular level, this might guide new avenues for treatment.

Insomnia is characterized by persistent difficulty initiating or maintaining sleep despite adequate sleep opportunity, along with associated daytime impairment^14^. An animal model of insomnia should recapitulate these characteristics and, in particular, display decreased ability to sleep despite environmental circumstances that normally promote sleep. Rodent models of insomnia generally involve perturbations such as stress or fear conditioning to activate arousal systems^15^, perhaps informative about acute insomnia (stress precipitated sleep loss), but less representative of chronic insomnia (conditioned sleeplessness). Neuro-imaging, EEG, and genetic work in humans lack the granularity to determine molecular mechanisms involved in onset and treatment of insomnia at a causal level. In contrast, short-sleeping *Drosophila* mutants are compelling models of chronic insomnia: reductions in sleep seen in numerous single gene mutants are primarily due to severely decreased sleep bout length, indicating that flies can initiate but not maintain sleep^16–19^. It is unlikely that these short sleepers simply do not need sleep, as mutants exhibit shortened lifespan and/or memory deficits^16–21^. In addition, a fly line generated by laboratory selection for insomnia-like traits^22^ shares many features of human insomnia, including reduced sleep time and consolidation, along with shortened lifespan and learning deficits. These fly models might therefore serve an important role in studying insomnia etiology and treatment.

Sleep quantity and quality also decrease with aging across species, including humans^23–25^. Moreover, recent work suggests a bidirectional relationship between sleep and Alzheimer’s disease (AD) pathology, where accumulation of the protein β-amyloid (Aβ) worsens sleep while poor sleep accelerates Aβ accumulation^26–28^. Indeed, in flies, Aβ accumulation in the brain leads to reduced and fragmented sleep^29^ and shortened lifespan^29,30^. While hypnotic use is associated with increased morbidity/mortality in individuals with AD^31^, behavioral therapies show promise for improving sleep^32–35^. Here, using principles of human behavioral sleep therapies in *Drosophila*, we developed a behavioral paradigm that markedly improves sleep quality in fly models of insomnia. We applied this approach to an AD model and found that sleep impairments due to Aβ are reversible with behavioral sleep modification; animals with improved sleep also show lifespan extension. Our findings demonstrate efficacy of behavioral sleep therapy in an experimentally-tractable system, establishing a new model to investigate the neurobiological basis of CBT-I.

## Methods

### Fly Strains

*Iso*^31^, *sleepless*^P1^, *redeye, period*^01^, *fumin*, and *cry*^02^ flies were obtained from A. Sehgal. *UAS:AβArctic* and *wide awake* were obtained from M. Wu. All of these lines were outcrossed at least 5x into the *iso*^31^ background. Canton S were obtained from E. Kravitz. Elav-Gal4 (#458) and *glass*^3^ (#508) were obtained from the Bloomington Drosophila Stock Center. Flies were maintained on standard yeast/cornmeal-based medium at 25 degrees on a 12hr:12hr LD cycle.

### Sleep Analysis

Male and female flies were collected at 1-3 days old and aged in group housing, and flipped onto new food every 3-4 days. Flies aged 5-8 days were loaded into glass tubes containing 5% sucrose and 2% agar. Locomotor activity was monitored using the Drosophila Activity Monitoring (DAM) system (Trikinetics, Waltham MA). Activity was measured in 1 min bins and sleep was defined as 5 minutes of consolidated inactivity^36^. Data was processed using PySolo software^37^. Sleep latency (SL) was determined by time (minutes) until first sleep episode following lights off. Wake after sleep onset (WASO) was calculated as the minutes of wake after initiation of the first sleep episode until end of the dark period. Activity index was calculated as the average number of beam breaks per minute of wake time. For all experiments, the first day of data following loading was discarded. Male flies were used for all experiments unless otherwise specified.

### Dark Time Extension

Five to eight day old flies were loaded into incubators and 2 days of baseline data were collected at 12:12 LD (9AM-9PM) cycles to compare populations at baseline. On day 3, light schedules either remained at 12:12 LD or shifted to a 10:14 LD or 8:16 LD cycle. Sleep data was collected for 4 additional days. Under 10:14 LD, incubators were dark from 8PM-10AM, while under 8:16 LD, incubators were dark from 7PM-11AM. Day 4-5 of data collection was used for analysis.

### Dark Time Restriction

Five to eight day old flies were loaded into incubators and 2 days of baseline data were collected at 12:12 LD (9AM-9PM) cycles to compare populations at baseline. On day 3, light schedules changed to the following (dark hours in parentheses): 20:4 LD (1AM-5AM) for days 3-4, 18:6 LD (12AM-6AM) for days 5-6, 16:8 LD (11PM-7AM) for days 7-8, and 14:10 LD (10PM to 8AM) for days 9-10 (Figure 2A). The 2nd day of each new LD cycle was used for analysis.

To evaluate the effects of tapering dark time, light schedules were changed to 18:6, 16:8 or 14:10 LD conditions, or the tapered restriction schedule above. 18:6 LD was compared to the tapered condition on Day 6, 16:8 LD was compared on day 8, and 14:10 LD was compared on day 10.

### Temperature Change

Five to eight day old flies previously entrained to 12:12LD conditions were loaded into incubators. Two days of baseline data were collected under DD (constant dark) conditions at 26°C to compare populations at baseline. On day 3, temperatures were reduced to 18°C during the following periods (otherwise at 26°C): 1AM-5AM for days 3-4, 12AM-6AM for days 5-6, 11PM-7AM for days 7-8, and 10PM to 8AM for days 9-10. The 2nd day of each new temperature cycle was used for analysis.

### Aging

Male and female flies were collected at 1-3 days old and group housed at a density of approximately 10 male and 10 female flies per vial. Flies were maintained on a dextrose-based food mixture, containing 11.7% (wt/vol) dextrose, 0.6% corn meal, and 0.3% yeast, and transferred to fresh food every 3-4 days. If fly density in vials became <10 flies, vials were combined to maintain original density. Flies were assayed for sleep and egg laying behaviors at 53 days post-eclosion.

### Egg Laying Assay

Egg laying assays were performed in 60 mm Petri dishes. Dishes were first filled with 8 ml molten dextrose-based food which was allowed to cool and solidify. Dishes were visually examined to ensure that the surface was smooth. Twenty aged female flies were placed upon a dextrose dish in an embryo collection cage (Genesee Scientific, cat#: 59-100). Dishes were replaced after 24 hours, and 3 consecutive days were averaged for each replicate experiment.

### Longevity Assay

Ten replicate vials, each containing 10 male and 10 female flies, were established for each condition. Flies were transferred to fresh dextrose-based food vials every 2-3 days, at which time dead flies were removed and recorded. Assays were conducted at least twice per genotype.

### Statistical Analysis and Data Reproducibility

Analysis was done using Prism (GraphPad Software). ANOVA with Tukey’s test was used in Figure 1C-H; Figure 2F-K; Figure 3B-D, F-H; Figure 4F-H, J-K; Figure 5B,D-I; Supplementary Figure 1A-J; Supplementary Figure 2A-I, L; Supplementary Figure 3A-I; Supplementary Figure 4B,G; and Supplementary Figure 5A. Student’s t-test was used in Figure 3J-L; Figure 4B-D, I, N-O; Supplementary Figure 1K-O; Supplementary Figure 2J-K; Supplementary Figure 3J-L; and Supplementary Figure 4C-F. Kolmogorov-Smirnov test was used in Figure 5A. Log-rank test was used in Figure 5J and Supplementary Figure 5B. For significance: *p≤0.05; **p<0.01; ***p<0.001. Each experiment was generated from a minimum of 3 independent biological replicates.

**Figure 1.**
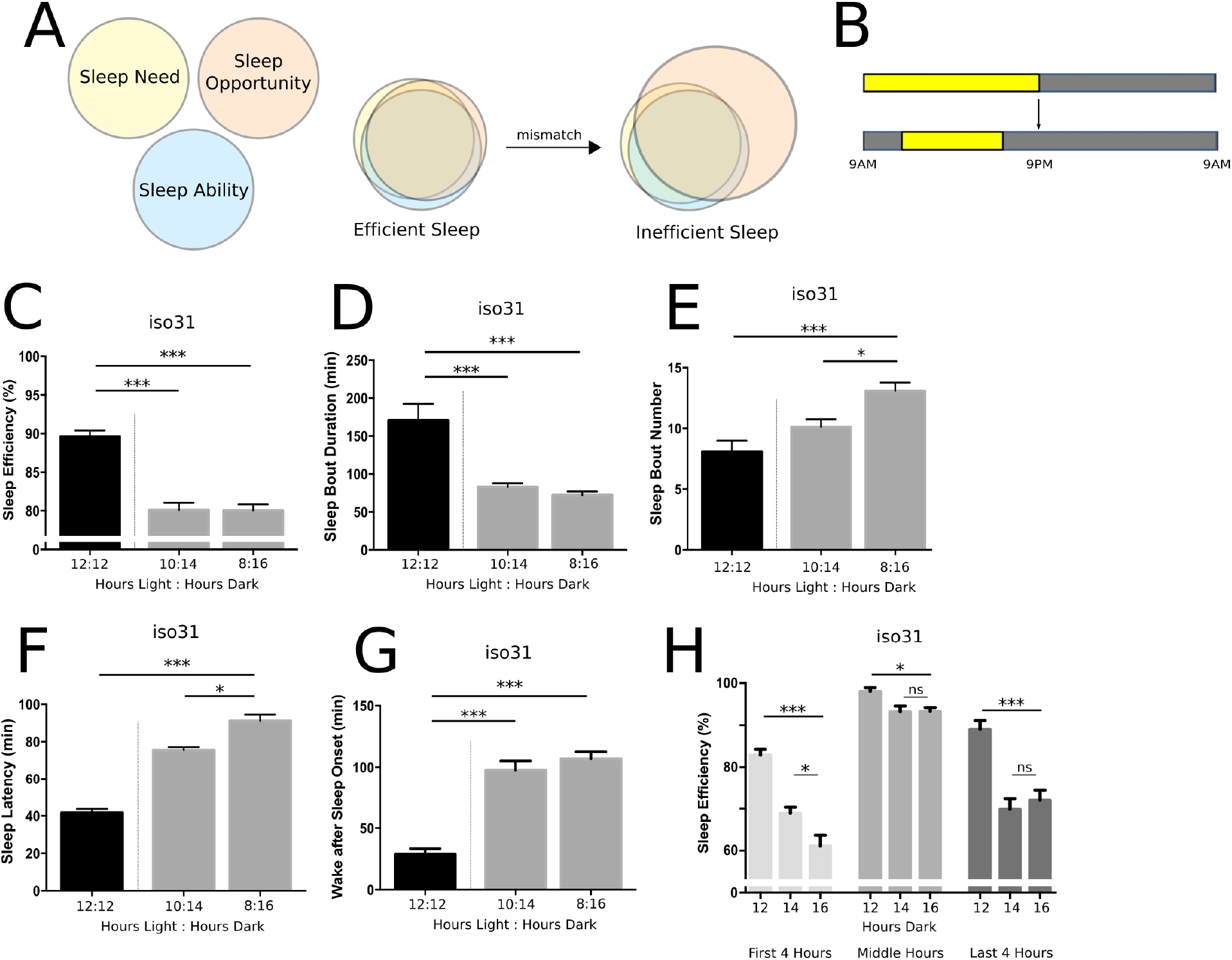
Sleep opportunity extension impairs sleep quality in *Drosophila*. (A) Schema of hypothesis of sleep degradation with mismatch of sleep opportunity and sleep ability. (B) Diagram of experimental extension of dark time from 12 hours (12:12 LD) to 16 hours (8:16 LD). Quantification of sleep efficiency (C), sleep bout duration (D), sleep bout number (E), sleep latency (F), and wake after sleep onset (G) following 3 nights of extended dark time in wild type iso^31^ flies (n = 48 flies). (H) Analysis of sleep efficiency based on time of dark period. For all figures, error bars represent SEM; *p < 0.05, **p < 0.01, ***p < 0.001.

**Figure 2.**
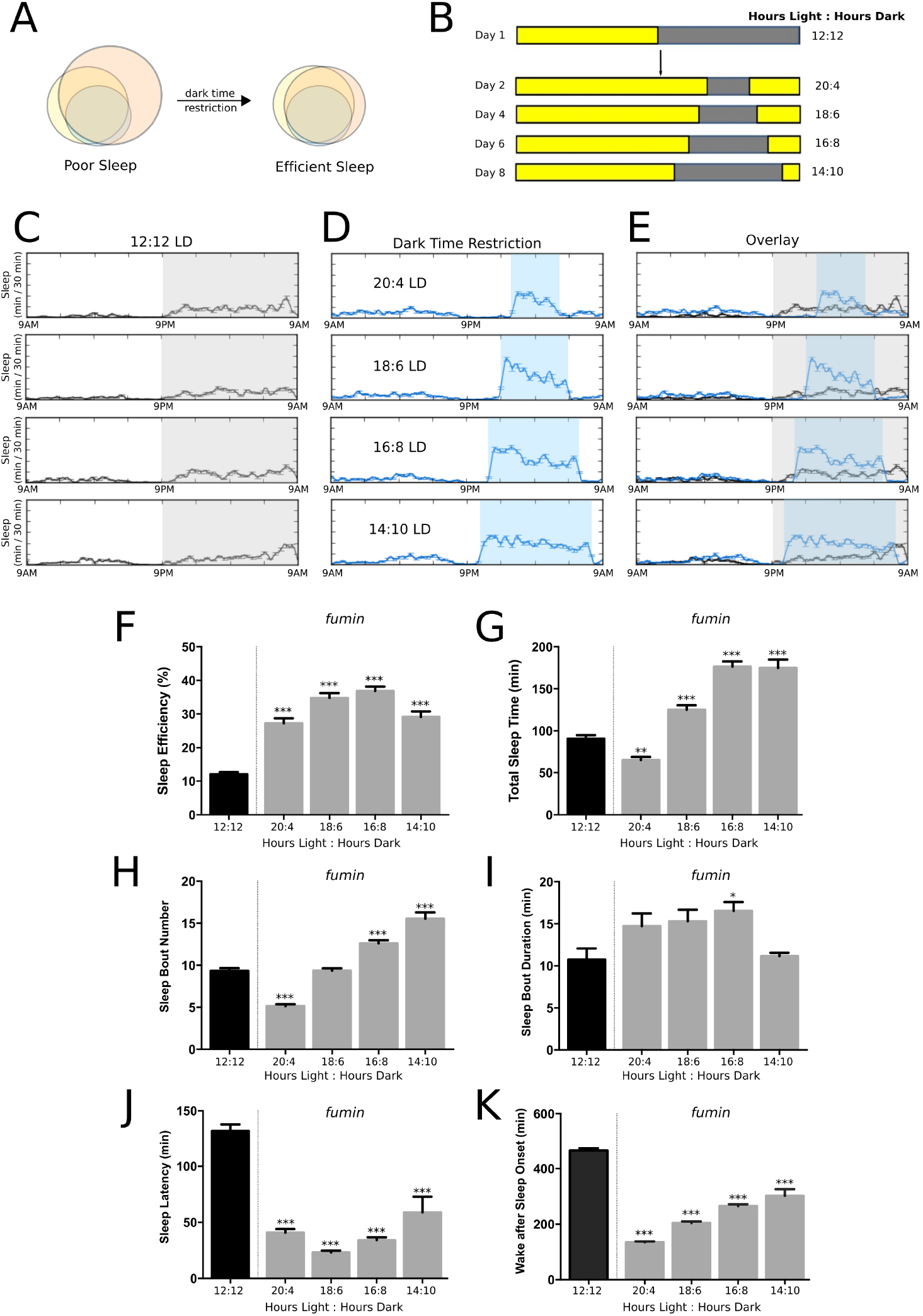
Sleep opportunity restriction enhances sleep in *fumin* mutants. (A) Schema of hypothesis that sleep compression aligns sleep opportunity and sleep ability, leading to efficient sleep. (B) Diagram of experimental protocol for restriction of dark time. (C-E) Representative sleep traces of *fumin* mutants under 12:12 LD conditions (C, gray shading indicates dark phase), sleep restriction protocol (D, blue shading indicates dark phase), and both plots overlaid (E). (F-K) Quantification of sleep measures with restriction of sleep opportunity in *fumin* mutants (n = 551 flies for 12:12 LD; n = 192 for 20:4 LD; n=204 for 18:6 LD; n=199 for 16:8 LD; and n=55 for 14:10 LD).

**Figure 3.**
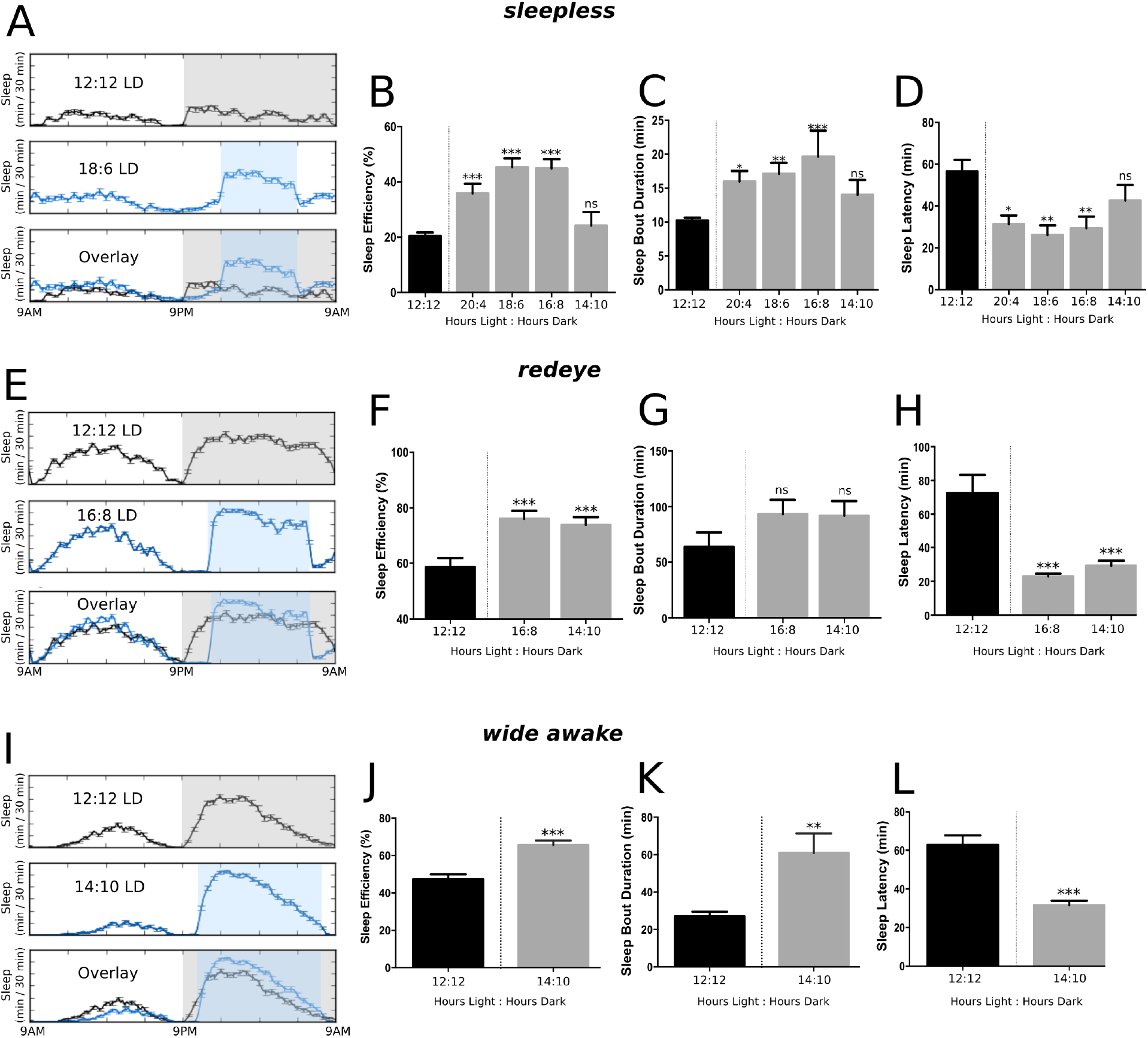
Sleep opportunity restriction improves sleep in multiple short-sleeping mutants. Representative sleep traces under 12:12 LD conditions (top panel, gray shading indicates dark phase), compressed sleep opportunity (middle panel, blue shading indicates dark phase) and overlaid plots (bottom panel) for *sleepless* (A), *redeye* (E), and *wide awake* (I) mutants. Quantification of sleep efficiency (B,F,J), sleep bout duration (C,G,K), and sleep latency (D,H,L) for each genotype (*sleepless*: n = 210 for 12:12 LD, n = 64 for 20:4 LD, n = 69 for 18:6 LD, n = 68 for 16:8 LD, and n = 33 for 14:10 LD; *redeye*: n = 63 for 12:12 LD, n = 60 for 16:8 LD, n = 58 for 14:10 LD; *wide awake*: n = 62 for 12:12 LD, n = 62 for 14:10 LD)

**Figure 4.**
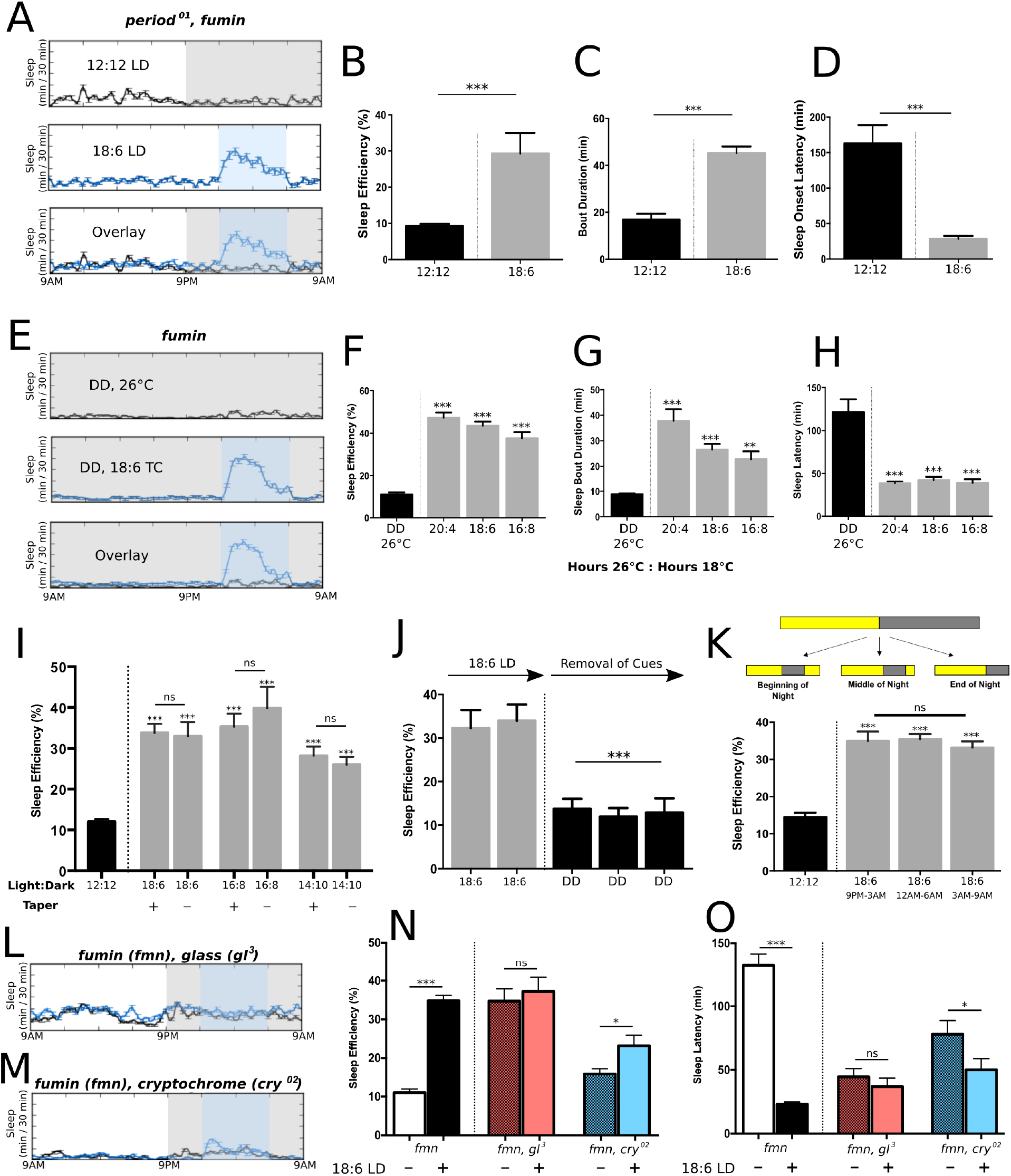
Response to sleep restriction requires ongoing environmental cues. (A) Representative sleep traces under 12:12 LD conditions (top panel, gray shading indicates dark phase), compressed sleep opportunity (middle panel, blue shading indicates dark phase) and overlaid plots (bottom panel) for *per*^01^; *fumin* mutants. Quantification of sleep efficiency (B), bout duration (C), and sleep latency (D) in *per*^01^; *fumin* mutants (n = 61 for 12:12 LD, n = 62 for 18:6 lD). (E) Representative sleep traces in *fumin* mutants under constant dark (DD) conditions (top panel, gray indicates 26°C) or with compressed sleep opportunity using temperature change (TC; middle panel, blue indicates 18°C; n = 144 for DD, n = 62 for 20:4 TC, n = 56 for 18:6 TC, n = 28 for 16:8 TC). Quantification of sleep efficiency (F), bout duration (G), and sleep latency (H) in *fumin* mutants under DD conditions with sleep opportunity restriction using temperature changes. (I) Sleep efficiency in *fumin* mutants with sleep restriction via tapered protocol versus sleep restriction initiated with the indicated dark period (n = 54,33 for 18:6 LD, n = 25,24 for 16:8 LD, n = 54,54 for 14:10 LD). (J) Sleep efficiency in *fumin* mutants under 18:6 LD conditions and after removal of environmental cues (n = 32). (K) Sleep efficiency in *fumin* mutants with 18:6 LD dark period restriction occurring at different times of night (n = 53 for 9p-3a, n = 172 for 12a-6a, n = 105 for 3a-9a). (L-O) Sleep opportunity restriction in light processing mutants. Overlaid sleep traces of *fumin;glass*^3^ (L) and *fumin;cry*^02^ (M). Black traces indicate 12:12 LD (gray shading indicates dark period); blue traces indicate sleep restriction (blue shading indicates dark period). Quantification of sleep efficiency (N) and sleep latency (O; n = 48 for *fumin;glass*^3^, n = 54 for *fumin;cry*^02^).

**Figure 5.**
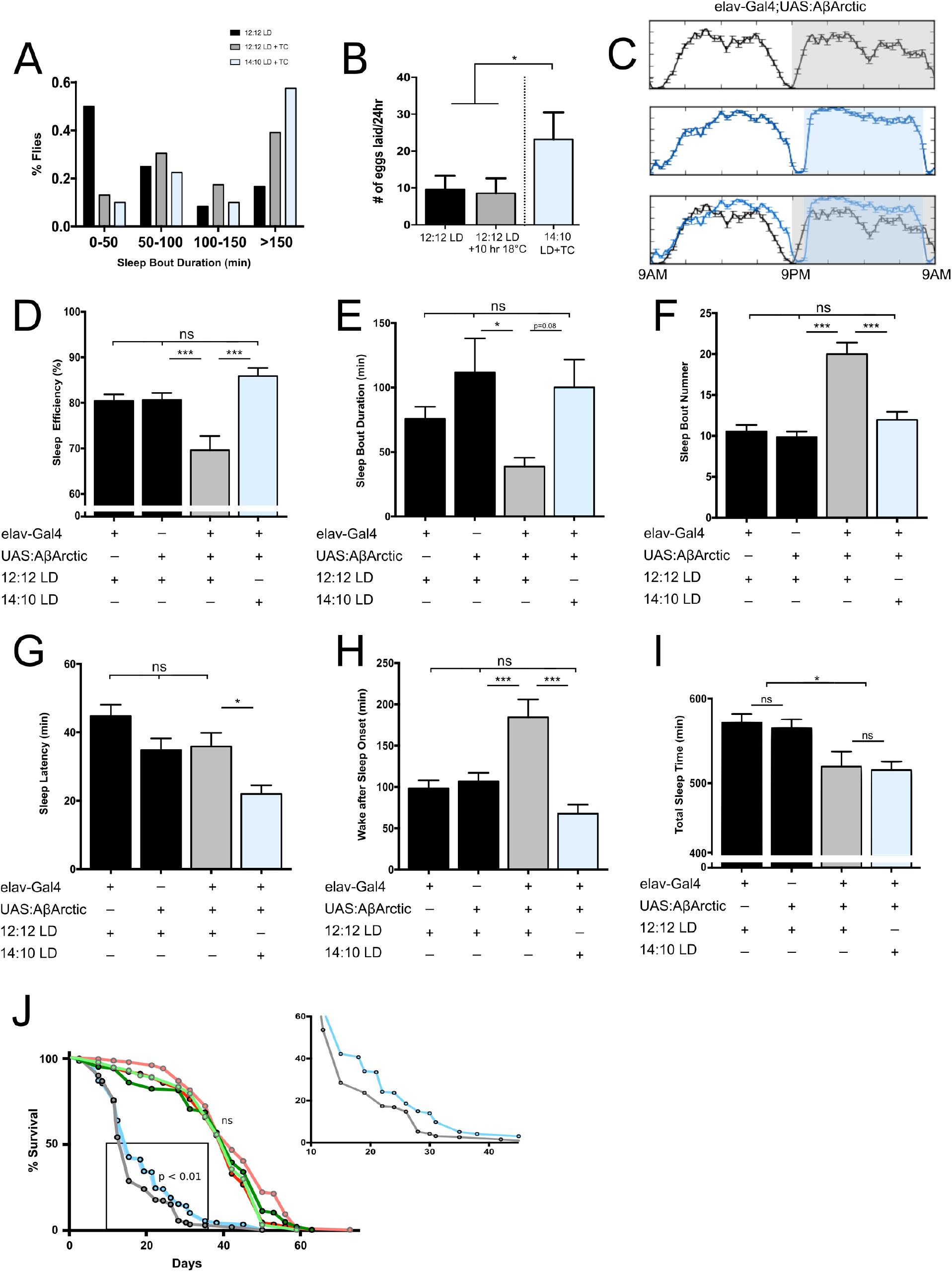
Sleep opportunity restriction improves sleep degradation associated with aging and enhanced Aβ burden. (A) Histogram of sleep bout lengths of aged flies (53 days old) under 12:12 LD (black bars, n = 75), 12:12 LD+TC (26°C:18°C, gray bars, n = 78), or 14:10 LD+TC conditions (blue bars, n = 77). (B) Number of eggs laid by aged female flies under 12:12 LD, 12:12 Ld plus 10 hours of low temperature during the light phase, or 14:10 LD+TC conditions (n = 100). (C) Representative sleep traces in flies with pan-neuronal overexpression of AβArctic under 12:12 LD conditions (top panel; gray shading indicates dark phase), compressed sleep opportunity (middle panel; blue shading indicates dark phase) and overlaid plots (bottom panel). (D-I) Quantification of sleep measures for *elav-Gal4/+* (n = 32), *UAS-AβArctic/+* (n = 32), and *elav-Gal4/UAS-AβArctic* (n = 32) flies under 12:12 LD or 14:10 LD conditions. (J) Survival curves with pan-neuronal overexpression of AβArctic or genetic controls under 12:12 LD or 14:10 LD conditions (n=100 males for each condition; *elav-Gal4/+:* 12:12 LD (light green) and 14:10 LD (dark green); *UAS:AβArctic/+:* 12:12 LD (light red) and 14:10 LD (dark red); *elav-Gal4/UAS:AβArctic:* 12:12 LD (gray) and 14:10 LD restriction (blue). Inset shows enlarged survival curves of *elav-Gal4/UAS:AβArctic* flies under each condition.

Samples were allocated based on genotype or experimental manipulation and statistics performed on aggregated data.

## Results

### Sleep opportunity extension impairs sleep quality in *Drosophila*

In aiming to model human behavioral sleep interventions in *Drosophila*, we first asked whether mismatch of sleep opportunity and ability degrades sleep quality in fruit flies (Fig. 1A), as it does it humans. Darkness is a powerful sleep-promoting cue in humans and *Drosophila*, and wild type flies raised on a 12hr:12hr light:dark (LD) cycle exhibit high sleep efficiency (sleep time divided by total sleep opportunity) over the dark period^38,39^. To control sleep timing and experimentally expand sleep opportunity, we examined sleep in wild type flies (*iso*^31^) following extension of the dark period from a baseline of 12 hours to 14 or 16 hours (Fig. 1B). This manipulation significantly decreased sleep efficiency, and increased sleep fragmentation as evidenced by shorter, more frequent sleep bouts (Fig. 1C-E). A similar effect on sleep following dark period extension was observed across multiple wild type strains (Supplementary Fig. 1A-C, 1F-H). In the clinical setting, measurement of sleep latency (SL) and time of wake after sleep onset (the cumulative wake duration during dark after the first sleep episode; WASO) are used to measure severity of sleep deficits^40^. We observed prolonged SL and increased WASO with dark period extension (Fig. 1F,G, Supplementary Fig. 1D-E, 1I-J). With sleep opportunity extension, flies specifically showed a large reduction in sleep efficiency during the first and last 4 hours of the night; a small decrease was observed in the middle hours of the night, but sleep efficiency remained over 90% (Fig. 1H). Thus, sleep degradation emerges primarily from loss of sleep quality at the beginning and end of the night. To test the role of the circadian clock in impaired sleep quality following sleep extension, we examined the *period* null mutant *per*^01^.^41^ While sleep efficiency was already low in these flies due to arrhythmicity (Supplementary Fig. 1K), sleep opportunity extension resulted in sleep fragmentation, prolonged SL, and increased WASO (Supplementary Fig. 1L-O), indicating that the response to sleep extension was not simply due to a mismatch in circadian timing. Together, these results suggest that, as in humans, flies cannot maintain efficient sleep when given an overabundance of sleep opportunity.

### Sleep opportunity restriction enhances sleep in short-sleeping mutants

If sleep extension results in analogous behavioral responses in humans and flies, can sleep opportunity restriction potentiate sleep efficiency in *Drosophila* short-sleeping mutants, as it does in humans with insomnia (Fig. 2A,B)? We first examined *fumin* mutants, which lack a functional dopamine transporter and sleep ~200-300 minutes per day, representing a 70-80% reduction from wild type levels (Fig. 2C)^42^. In humans with insomnia undergoing CBT-I, the initial amount of sleep restriction is determined based on an individual’s total sleep time (TST); a titration procedure is then used to increase sleep opportunity as sleep is consolidated and becomes more efficient^43^. Applying this approach to *fumin* mutants, sleep time was compressed by initially contracting dark time to 4 hours, followed by titration of sleep opportunity by expanding the dark period by 2 hours every other day (Fig. 2B). Using this paradigm, we observed a threefold increase in sleep efficiency, with maximal improvement at 6-8 hours sleep opportunity (Fig 2C-F).

Enhanced sleep efficiency was not simply a function of comparing sleep within a compressed dark period to the entire 12 hours of dark: sleep efficiency in the restricted condition was also elevated in comparison to non-restricted flies (LD 12:12) during the same smaller time window or the equivalent number of hours following start of the dark period (Supplementary Fig. 2A-D). Interestingly, TST with compression of the dark period was increased above 12:12 LD conditions (Fig. 2G), despite reduced opportunity. The enhancement in sleep efficiency and TST was driven primarily by an increase in the number of sleep bouts initiated with sleep opportunity restriction (Fig. 2H). With only 6 hours of sleep opportunity, *fumin* flies initiated the same number of bouts that normally occurred during the entire 12 hours of sleep under LD 12:12 conditions (Fig. 2H); indeed, comparison of the same 6 hour dark period under LD 12:12 and LD 18:6 conditions revealed that restricted flies exhibit significantly more sleep bouts during this time (Supplementary Fig. 2E). Moreover, given 8 hours of sleep opportunity (LD 16:8), *fumin* flies initiate even more sleep bouts than within the entire 12 hour period under non-sleep restricted conditions (Fig. 2H). In contrast, only a modest increase in sleep bout duration was observed with compression of sleep opportunity (Fig. 2I), indicating that *fumin* flies initiate more bouts with matching of sleep opportunity and ability, without much change in sleep maintenance. Both SL and WASO were significantly decreased under all dark time-restricted conditions (Fig. 2J,K), further indication of improved sleep quality.

Importantly, similar restriction of sleep opportunity in wild type flies did not increase sleep efficiency, perhaps because of a ceiling effect (baseline ~90%, Supplementary Fig. 2F). While there was a trend towards more consolidated sleep in wild type flies with a compressed dark period (Supplementary Fig. 2G,H), this occurred in the setting of daytime rebound sleep (Supplementary Fig. 2I). These results indicate that, as would be expected, restricting sleep opportunity in efficient-sleeping wild type flies induces a state of sleep deprivation and associated homeostatic compensation. In contrast, restriction of sleep opportunity to as little as 6 hours in *fumin* mutants did not induce subsequent daytime rebound sleep or a change to daytime activity (Supplementary Fig. 2J-L), suggesting that sleep opportunity and ability become better matched with sleep restriction.

To test whether enhanced sleep following sleep opportunity restriction is specific to *fumin* mutants, we next examined this paradigm in other mutants with distinct genetic lesions underlying the short-sleep phenotype: *sleepless, redeye*, and *wide* awake^17,44,45^. The dark period was restricted based on average TST for each mutant under 12:12 LD cycles. As with *fumin* flies, we found that restriction of sleep opportunity in each mutant increased sleep efficiency and consolidation, and reduced SL and WASO (Fig. 3, Supplementary Fig. 3). TST was largely unchanged with sleep compression (Supplementary Fig. 3), suggesting that the primary improvement is in sleep quality, consistent with CBT-I findings in humans^46–48^. These results demonstrate that behavioral sleep modification can be applied across a variety of short-sleep etiologies, and indicate there is a ceiling beyond which sleep cannot be improved (i.e., sleep mutants cannot be fully restored to wild type sleep levels). Together, our data establish a paradigm for behavioral sleep modification in flies, and suggest that sleep ability is plastic in *Drosophila* short-sleeping mutants.

### Response to sleep restriction requires ongoing environmental cues

All of the short-sleeping mutants we examined have normal molecular and behavioral rhythms under 12:12 LD conditions^17,42,44,45^, but aberrant light cycles affect function of the molecular clock. To determine whether enhanced sleep following sleep opportunity restriction is clock-dependent, we generated *per*^01^;*fumin* double mutants that lack a functional molecular clock in addition to exhibiting a short-sleep phenotype. With dark period compression, we observed that increased sleep efficiency and consolidation persist, indicating that sleep restriction is clock-independent (Fig. 4A-D). This finding is consistent with previous work showing the circadian clock requires at least 8 hours of dark time for robust cycling^49^, such that 4 or 6 hours of dark, in which enhanced sleep was observed in *fumin*, likely disrupted clock function.

We next asked if sleep restriction is specific to dark as a sleep-permissive cue. Cool temperatures are also sleep-permissive in both humans and flies^50,51^, and under constant dark conditions, flies exhibit consolidated sleep at subjective night with lower temperature^52^. Using temperature changes (TC) from warm (26°C) to cool (18°C) under constant darkness (DD), we compared DD conditions at 26°C to restricted periods of low temperature in *fumin* flies. Restriction of sleep opportunity with temperature, like darkness, caused increased sleep efficiency, increased bout length, and decreased SL, indicating that enhanced sleep with sleep restriction is not specific to light/dark inputs (Fig. 4E-H). Restriction of sleep opportunity with coincident darkness and low temperature yielded similar increased sleep efficiency to dark or low temperature alone (Supplementary Fig. 4A,B), suggesting that either cue is sufficient for the maximum sleep improvement in *fumin* mutants.

How do other features of behavioral sleep modification in humans function in our fly model? First, in humans, behavioral sleep modification initiates with the greatest restriction of sleep opportunity and the goal of enhancing sleep drive/stabilizing sleep ability. This is followed by increased periods of sleep opportunity (titration) that would not have yielded efficient sleep at the outset. To test whether the titration paradigm is necessary in flies, we examined gradual extension of the dark period from 4 to 10 hours in comparison to direct initiation of sleep opportunity restriction at either 6, 8, or 10 hours in *fumin* mutants. We found that enhanced sleep efficiency and sleep consolidation was similar whether tapered from 4 hours or restricted directly to 6, 8, or 10 hours (Fig. 4I, Supplementary Fig. 4C,D). Second, improved sleep with sleep restriction in humans can take days to manifest, as sleep drive builds. We found in *fumin* mutants that the first day of sleep opportunity restriction (whether 4 or 6 hours) did not induce a maximal improvement in sleep efficiency or SL compared to 12:12 LD conditions; improvement reliably maximized by day 3 of restriction (Supplementary Fig. 4E,F), suggesting that homeostatic sleep drive has to build over time. Third, adherence to the components of CBT-I, including sleep restriction, is strongly related to treatment outcome^53^. We asked whether enhanced sleep with dark period compression persists with termination of sleep restriction. We restricted sleep opportunity in *fumin* mutants to 6 hours (18:6 LD), and then shifted to constant dark (DD) to test if increased sleep efficiency continues in the absence of all light cues. With this manipulation, we found a regression of sleep efficiency back to baseline (Fig. 4J). Finally, humans with insomnia who undergo behavioral sleep modification might restrict sleep from the beginning of the night, end, or both depending on patient preference. We initially modeled *Drosophila* sleep restriction by limiting sleep opportunity from both start and end of the night (e.g., Zeitgeber Time (ZT) 15-21 for 6 hours of restriction). To test whether this behavioral paradigm depends on timing of sleep restriction or only total amount, we limited sleep opportunity to either the first 6 (ZT 12-18) or last 6 (ZT 18-24) hours of the subjective night. We observed no significant difference in sleep efficiency or SL across these conditions (Fig. 4K, Supplementary Fig. 4G), indicating that the amount of sleep opportunity, not the timing, determines response.

Our results suggest that blocking sensory processing of LD cues should occlude the response to sleep restriction with dark period compression. Flies process light through canonical visual pathways as well as other light sensors such as CRYPTOCHROME (CRY)^54^; genetic disruption of both of these pathways renders *Drosophila* insensitive to LD cycling and behavioral arrhythmicity^54,55^. We generated *glass;fumin* double mutants, which lack all functional eye components and are short-sleepers. These double mutants exhibited no change in sleep efficiency or SL with dark period restriction (Fig. 4M-O), indicating that a functional eye is necessary for induction of sleep restriction. CRY is a UV- and blue light-sensitive protein that communicates light information to the circadian system^56–58^. To test whether CRY plays a role in the response to sleep opportunity restriction, we generated *cry*^02^:*fumin* double mutants. These flies exhibited an attenuated response to sleep restriction (Fig. 4L,N,O), suggesting that maximal increases in sleep efficiency with restriction of the dark period utilize multiple light-processing systems. Together, these data demonstrate that sleep restriction has a direct reliance on environmental cues to produce its effect regardless of prior experience, and that sleep opportunity restriction in flies does not cause a long-lasting change in the absence of these cues.

### Sleep restriction improves sleep in aging and Alzheimer’s disease models

Aging is associated with increased sleep fragmentation in Drosophila^59–61^ and humans^62^. We next investigated whether behavioral sleep modification through sleep restriction might improve sleep quality in aged flies. Behavioral response to light cues are weakened in aged flies, but sleep can be consolidated by adding coincident temperature cycles to 12:12 LD cycles^63^. To investigate whether aged flies further consolidate sleep with sleep opportunity restriction, we compared aged female flies (53 days post-eclosion) under 12:12 LD+TC (26°C:18°C) conditions to flies that were restricted to 10 hours dark and coincident low temperature. We chose 10 hours of sleep opportunity to match TST during the night at baseline. We observed an increase in sleep bout length in restricted flies, above that of 12:12 LD+TC alone, indicating a consolidation of sleep with restriction (Fig. 5A). No significant increase in sleep efficiency was observed above 12:12 LD+TC, likely due to a ceiling effect in sleep efficiency in aged flies (Supplementary Fig. 5A). To assess behavioral consequences of consolidating sleep in aged flies, we examined reproductive fitness following sleep opportunity restriction. Aged female flies normally exhibit a dramatic reduction in reproductive output^64^, and reproductive output is also impaired with sleep deprivation^65^. We tested whether such decrements are modifiable with improved sleep. We assessed egg laying behavior after 5 nights of sleep opportunity restriction in 53 day old mated female flies, and found that flies with improved sleep laid significantly more eggs in a 24-hour period than controls (Fig. 5B). This increase was not simply due to exposure to cool temperatures, as addition of an equivalent low temperature period during the day under 12:12 LD conditions was indistinguishable from control flies. These results raise the possibility of potential behavioral benefits to improved sleep in aged flies following restriction of sleep opportunity.

Sleep quality degrades with normal aging, but disruptions to sleep are also increasingly appreciated in neurodegenerative processes like Alzheimer’s disease (AD)^66^. Several models of AD have been established in *Drosophila*, including those based on expression of aggregating β-amyloid (Aβ) peptides^67,68^; Aβ accumulation results in decreased and fragmented sleep, while sleep deprivation increases Aβ burden^29^. We examined sleep following pan-neuronal expression of AβArctic, which carries a mutation to induce enhanced aggregation^67^. Consistent with previous work^29^, we observed reduced night TST and increased sleep fragmentation in 7-10 day old male flies with pan-neuronal AβArctic expression under 12:12 LD cycles (Fig. 5C,E,F,I). Sleep was less efficient, due to a reduction in sleep bout duration and increase in number of sleep bouts (Fig. 5D-F); WASO was likewise increased with pan-neuronal AβArctic expression, though SL was unaffected (Fig. 5G,H). We next examined whether sleep degradation related to Aβ accumulation is reversible with sleep opportunity restriction using dark period compression. We found this manipulation restored sleep efficiency, sleep bout length, and number of sleep bouts back to control levels (Fig. 5D-F); WASO was also normalized, and SL was shortened (Fig. 5G,H). TST was unchanged with restriction of sleep opportunity, but flies were able to achieve the same amount of sleep in the compressed window (Fig. 5I). Thus, manipulation of environmental cues is sufficient to improve sleep despite pan-neuronal Aβ aggregation.

Does enhancement of sleep quality in this model of AD have other beneficial effects? AβArctic flies exhibit severly curtailed lifespan^29^, so we tested whether correcting sleep can affect longevity. Comparing flies expressing AβArctic pan-neuronally under either 12:12 LD or dark-restricted conditions, we found that sleep opportunity restriction was associated with a significant extension of lifespan in both males and females (Fig. 5J, Supplementary Fig. 5B). This longevity extension was not due to changes in the LD cycle, as genetic controls showed no alteration in longevity with sleep opportunity restriction. Taken together, these data suggest that behavioral sleep modification mitigates Aβ-related sleep disturbances and shortened lifespan.

## Discussion

CBT-I is the first-line treatment for insomnia, offering advantages over existing pharmacotherapies with regard to safety and durability of response^33^. However, CBT-I is limited by obstacles to broad implementation^11–13^. Research in *Drosophila* has yielded numerous insights into basic sleep neurobiology, and here, we have leveraged this system to develop a tractable experimental model of sleep restriction therapy for insomnia. We find that mismatch of sleep opportunity and ability degrades sleep quality in flies, as in humans. Surprisingly, compression of sleep opportunity in short-sleeping genetic mutants improves sleep efficiency along with multiple measures of sleep quality. We apply this paradigm to normal aging and neurodegeneration, both of which are associated with impaired sleep, and find that behavioral sleep modification restores sleep quality and extends lifespan. These data establish a new platform for deciphering mechanistic principles of a behavioral sleep therapy that improves sleep across species.

Previous work has argued that short-sleeping flies are a compelling model for studying human insomnia^22,69^. Single gene mutants such as those tested here^17,42,44,45^, as well as a line generated by laboratory selection over many generations^22^, recapitulate central features of human insomnia: reduced sleep time, increased sleep latency and sleep fragmentation, and daytime impairments. The conserved response in flies to both sleep opportunity extension and restriction provides further support for the idea that this organism can serve as a valid model for insomnia. Human evidence is consistent with a genetic component to insomnia^70,71^, and while this disease is likely multigenic in nature^72^, highly penetrant single gene mutations are important for studying disorder mechanisms and treatment approaches. The fact that genetically-distinct *Drosophila* sleep mutants all respond to the sleep compression paradigm suggests these lesions ultimately converge on a shared physiological, and perhaps cellular, endpoint. Our data also indicate that sleep ability is plastic: optimizing environmental conditions can enhance sleep efficiency (and even total sleep time in *fumin* flies; Fig. 2G) despite fixed genetic mutations, suggesting biological determinants of sleep are highly mutable. This idea is conceptually informative for humans with insomnia, and provides empirical evidence for focusing on mechanisms of sleep opportunity restriction as the core insomnia treatment modality. Indeed, the Spielman model for insomnia (also known as the 3P model) identifies predisposing (e.g., genetic) and precipitating factors (e.g., acute stressor) that lead to acute insomnia, with perpetuating factors (e.g., sleep extension) that shift acute insomnia to chronic^73,74^. This model has served as the basis for using sleep restriction in humans to target sleep extension (a perpetuating factor). Our results raise the possibility that sleep restriction also targets predisposing genetic factors, by better matching intrinsic sleep ability with opportunity. In other words, humans with a genetic predisposition to insomnia might be sleep “over-extended” even if sleep opportunity appears normal; restriction of sleep opportunity would therefore increase sleep efficiency and perhaps potentiate sleep ability.

Poor sleep has long been appreciated as a comorbidity of aging and neurodegeneration^25,28^, but more recently identified as a potential modifiable risk factor for neurodegenerative disease progression^26–28^. In flies, pharmacologic and genetic approaches to improve sleep have been shown to ameliorate memory deficits in an Alzheimer’s disease model^75^; similarly, altering the sleep-Aβ interaction by modulating neuronal excitability with a pharmacotherapy prolongs lifespan in Aβ-expressing flies^29^. We find that increased sleep efficiency through compression of sleep opportunity is alone sufficient to extend lifespan in Aβ-expressing flies. Intriguingly, this work suggests that behavioral approaches to treating insomnia could slow progression of disease, consistent with evidence in humans demonstrating that CBT-I in older adults with mild cognitive impairment improves cognitive function^76^.

Most pharmacological treatments in psychiatry are based on drugs discovered serendipitously over a half century ago^77^. In recent years, significant advances in treating mental illness have been behavioral interventions^78^, yet little is known regarding the mechanistic basis of such interventions. How can behavioral therapies be studied at a molecular level? This fly model of behavioral sleep modification can be used to generate such granular insights. Our initial results demonstrate that therapeutic sleep restriction does not require a functional molecular clock, and that manipulating light:dark cycles to enhance sleep drive requires canonical light sensory pathways. Future work will use this model to define the neural circuits required for, and molecular changes occurring with, sleep restriction, with the goal of identifying new insomnia treatment targets that are conceptually based on the established efficacy of CBT-I.

## Acknowledgements

We thank Kyunghee Koh, Philip Gehrman, and members of the Kayser Lab for helpful discussions. This work was supported by NIH grants F30AG058409 (SJB), T32HL07953 (SJB), R21HD083628 (MLP), K24AG055602 (MLP), K08NS090461 (MSK), a Hearst Foundation Fellowship Award (SJB), and Burroughs Wellcome Career Award for Medical Scientists, March of Dimes Basil O’Connor Scholar Award, and Sloan Research Fellowship (MSK).

## References

1. Dement, W. & Pelayo, R. Public health impact and treatment of insomnia. Eur. Psychiatry 12, 31s–39s (1997).

2. Ozminkowski, R. J., Wang, S. & Walsh, J. K. The direct and indirect costs of untreated insomnia in adults in the United States. Sleep 30, 263–273 (2007).

3. Daley, M. et al. Insomnia and its relationship to health-care utilization, work absenteeism, productivity and accidents. Sleep Med. 10, 427–438 (2009).

4. Sarsour, K., Morin, C. M., Foley, K., Kalsekar, A. & Walsh, J. K. Association of insomnia severity and comorbid medical and psychiatric disorders in a health plan-based sample: Insomnia severity and comorbidities. Sleep Med. 11, 69–74 (2010).

5. Qaseem, A. et al. Management of chronic insomnia disorder in adults: A clinical practice guideline from the American college of physicians. Ann. Intern. Med. 165, 125–133 (2016).

6. Miller, C. B. et al. The evidence base of sleep restriction therapy for treating insomnia disorder. Sleep Medicine Reviews 18, 415–424 (2014).

7. Spielman, A. J. Assessment of insomnia. Clin. Psychol. Rev. 6, 11–25 (1986).

8. Morin, C. M., LeBlanc, M., Daley, M., Gregoire, J. P. & Mérette, C. Epidemiology of insomnia: Prevalence, self-help treatments, consultations, and determinants of help-seeking behaviors. Sleep Med. 7, 123–130 (2006).

9. Trauer, J. M., Qian, M. Y., Doyle, J. S., Rajaratnam, S. M. W. & Cunnington, D. Cognitive behavioral therapy for chronic insomnia: A systematic review and meta-analysis. Ann. Intern. Med. 163, 191–204 (2015).

10. Wu, J. Q., Appleman, E. R., Salazar, R. D. & Ong, J. C. Cognitive behavioral therapy for insomnia comorbid with psychiatric and medical conditions a meta-analysis. JAMA Intern. Med. 175, 1461–1472 (2015).

11. Sivertsen, B., Vedaa, Ø. & Nordgreen, T. The Future of Insomnia Treatment—the Challenge of Implementation. Sleep 36, 303–304 (2013).

12. Kathol, R. G. & Arnedt, J. T. Cognitive behavioral therapy for chronic insomnia: Confronting the challenges to implementation. Annals of Internal Medicine 165, 149–150 (2016).

13. Mitchell, M. D., Gehrman, P., Perlis, M. & Umscheid, C. A. Comparative effectiveness of cognitive behavioral therapy for insomnia: A systematic review. BMC Family Practice 13, (2012).

14. Roth, T. Insomnia: Definition, prevalence, etiology, and consequences. Journal of Clinical Sleep Medicine 3, (2007).

15. Toth, L. A. & Bhargava, P. Animal models of sleep disorders. Comp. Med. 63, 91–104 (2013).

16. Cirelli, C. et al. Reduced sleep in Drosophila Shaker mutants. Nature 434, 1087–1092 (2005).

17. Koh, K. et al. Identification of SLEEPLESS, a Sleep-Promoting Factor. Science (80-.). 321, 372–376 (2008).

18. Bushey, D., Huber, R., Tononi, G. & Cirelli, C. Drosophila Hyperkinetic mutants have reduced sleep and impaired memory. J. Neurosci. 27, 5384–5393 (2007).

19. Stavropoulos, N. & Young, M. W. Article insomniac and Cullin-3 Regulate Sleep and Wakefulness in Drosophila. Neuron 72, 964–976 (2011).

20. Bushey, D., Hughes, K. a, Tononi, G. & Cirelli, C. Sleep, aging, and lifespan in Drosophila. BMC Neurosci. 11, 56 (2010).

21. Zhang, S., Yin, Y., Lu, H. & Guo, A. Increased dopaminergic signaling impairs aversive olfactory memory retention in Drosophila. Biochem. Biophys. Res. Commun. 370, 82–86 (2008).

22. Seugnet, L. et al. Identifying Sleep Regulatory Genes Using a Drosophila Model of Insomnia. J. Neurosci. 29, 7148–7157 (2009).

23. Robertson, M. & Keene, A. C. Molecular Mechanisms of Age-Related Sleep Loss in the Fruit Fly - A Mini-Review. Gerontology 59, 334–339 (2013).

24. Hasan, S., Dauvilliers, Y., Mongrain, V., Franken, P. & Tafti, M. Age-related changes in sleep in inbred mice are genotype dependent. Neurobiol. Aging 33, (2012).

25. Mander, B. A., Winer, J. R. & Walker, M. P. Sleep and Human Aging. Neuron 94, 19–36 (2017).

26. Kang, J.-E. E. et al. Amyloid-B Dynamics Are Regulated by Orexin and the Sleep-Wake Cycle. Science (80-.). 326, 1005–1007 (2009).

27. Roh, J. H. et al. Disruption of the sleep-wake cycle and diurnal fluctuation of beta-amyloid in mice with Alzheimer’s disease pathology. Sci Transl Med 4, 150ra122 (2012).

28. Ju, Y.-E. S., Lucey, B. P. & Holtzman, D. M. Sleep and Alzheimer disease pathology—a bidirectional relationship. Nat. Rev. Neurol. 10, 115–119 (2014).

29. Tabuchi, M. et al. Sleep interacts with aβ to modulate intrinsic neuronal excitability. Curr. Biol. 25, 702–12 (2015).

30. Iijima, K. et al. Dissecting the pathological effects of human Aβ40 and Aβ42 in Drosophila: A potential model for Alzheimer’s disease. Proc. Natl. Acad. Sci. U. S. A. 101, 6623–6628 (2004).

31. Saarelainen, L. et al. Risk of death associated with new benzodiazepine use among persons with Alzheimer disease: A matched cohort study. Int J Geriatr Psychatry 33, 583–590 (2018).

32. Krystal, A. D. & Edinger, J. D. Sleep EEG predictors and correlates of the response to cognitive behavioral therapy for insomnia. Sleep 33, 669–77 (2010).

33. Riemann, D. & Perlis, M. L. The treatments of chronic insomnia: A review of benzodiazepine receptor agonists and psychological and behavioral therapies. Sleep Medicine Reviews 13, 205–214 (2009).

34. Ngandu, T. et al. A 2 year multidomain intervention of diet, exercise, cognitive training, and vascular risk monitoring versus control to prevent cognitive decline in at-risk elderly people (FINGER): A randomised controlled trial. Lancet 385, 2255–2263 (2015).

35. Kivipelto, M. et al. The Finnish Geriatric Intervention Study to Prevent Cognitive Impairment and Disability (FINGER): Study design and progress. Alzheimer’s Dement. 9, 657–665 (2013).

36. Gilestro, G. F. Video tracking and analysis of sleep in Drosophila melanogaster. Nat. Protoc. 7, 995–1007 (2012).

37. Gilestro, G. F. & Cirelli, C. PySolo: A complete suite for sleep analysis in Drosophila. Bioinformatics 25, 1466–1467 (2009).

38. Hendricks, J. C. et al. Rest in Drosophila Is a Sleep-like State. Neuron 25, 129–138 (2000).

39. Shaw, P. J., Cirelli, C., Greenspan, R. J. & Tononi, G. Correlates of Sleep and Waking in Drosophila melanogaster. 287, (2000).

40. Sateia, M. J., Doghramji, K., Hauri, P. J. & Morin, C. M. Evaluation of chronic insomnia. An American Academy of Sleep Medicine review. Sleep 23, 243–308 (2000).

41. Konopka, R. J. & Benzer, S. Clock mutants of Drosophila melanogaster. Proc. Natl. Acad. Sci. U. S. A. 68, 2112–6 (1971).

42. Kume, K., Kume, S., Park, S. K., Hirsh, J. & Jackson, F. R. Dopamine Is a Regulator of Arousal in the Fruit Fly. 25, 7377–7384 (2005).

43. Kyle, S. D. et al. Towards standardisation and improved understanding of sleep restriction therapy for insomnia disorder: A systematic examination of CBT-I trial content. Sleep Medicine Reviews 23, 83–88 (2015).

44. Shi, M., Yue, Z., Kuryatov, A., Lindstrom, J. M. & Sehgal, A. Identification of Redeye, a new sleep-regulating protein whose expression is modulated by sleep amount. 1–17 (2014). doi:10.7554/eLife.01473

45. Liu, S. et al. WIDE AWAKE mediates the circadian timing of sleep onset. Neuron 82, 151–166 (2014).

46. Edinger, J. D., Wohlgemuth, W. K., Radtke, R. a, Marsh, G. R. & Quillian, R. E. Cognitive behavioral therapy for treatment of chronic primary insomnia: a randomized controlled trial. JAMA 285, 1856–64 (2001).

47. Smith, M. T. et al. Comparative meta-analysis of pharmacotherapy and behavior therapy for persistent insomnia. American Journal of Psychiatry 159, 5–11 (2002).

48. Edinger, J. D., Wohlgemuth, W. K., Radtke, R. A., Coffman, C. J. & Carney, C. E. Dose-response effects of cognitive-behavioral insomnia therapy: A randomized clinical trial. Sleep 30, 203–212 (2007).

49. Qiu, J. & Hardin, P. E. per mRNA cycling is locked to lights-off under photoperiodic conditions that support circadian feedback loop function. Mol. Cell. Biol. 16, 4182–8 (1996).

50. Okamoto-Mizuno, K. & Mizuno, K. Effects of thermal environment on sleep and circadian rhythm. Journal of Physiological Anthropology 31, 1–9 (2012).

51. Kaneko, H. et al. Circadian rhythm of temperature preference and its neural control in Drosophila. Curr. Biol. 22, 1851–1857 (2012).

52. Ishimoto, H., Lark, A. & Kitamoto, T. Factors that differentially affect daytime and nighttime sleep in Drosophila melanogaster. Front. Neurol. FEB, 1–5 (2012).

53. Matthews, E. E., Arnedt, J. T., McCarthy, M. S., Cuddihy, L. J. & Aloia, M. S. Adherence to cognitive behavioral therapy for insomnia: A systematic review. Sleep Medicine Reviews 17, 453–464 (2013).

54. Yoshii, T., Hermann-Luibl, C. & Helfrich-Förster, C. Circadian light-input pathways in Drosophila. Communicative and Integrative Biology 9, (2016).

55. Helfrich-Förster, C., Winter, C., Hofbauer, A., Hall, J. C. & Stanewsky, R. The circadian clock of fruit flies is blind after elimination of all known photoreceptors. Neuron 30, 249–261 (2001).

56. Emery, P., So, W. V., Kaneko, M., Hall, J. C. & Rosbash, M. Cry, a Drosophila clock and light-regulated cryptochrome, is a major contributor to circadian rhythm resetting and photosensitivity. Cell 95, 669–679 (1998).

57. Emery, P. et al. Drosophila CRY is a deep brain circadian photoreceptor. Neuron 26, 493–504 (2000).

58. Agrawal, P. et al. Drosophila CRY Entrains Clocks in Body Tissues to Light and Maintains Passive Membrane Properties in a Non-clock Body Tissue Independent of Light. Curr. Biol. 27, 2431–2441.e3 (2017).

59. Koh, K., Evans, J. M., Hendricks, J. C. & Sehgal, A. A Drosophila model for age-associated changes in sleep:wake cycles. Proc. Natl. Acad. Sci. U. S. A. 103, 13843–13847 (2006).

60. Brown, M. K. et al. Aging induced endoplasmic reticulum stress alters sleep and sleep homeostasis. Neurobiol. Aging 35, 1431–1441 (2014).

61. Vienne, J., Spann, R., Guo, F. & Rosbash, M. Age-Related Reduction of Recovery Sleep and Arousal Threshold in Drosophila. Sleep 39, 1613–1624 (2016).

62. Pandi-Perumal, S. R. et al. Senescence, sleep, and circadian rhythms. Ageing Research Reviews 1, 559–604 (2002).

63. Luo, W. et al. Old flies have a robust central oscillator but weaker behavioral rhythms that can be improved by genetic and environmental manipulations. Aging Cell 11, 428–438 (2012).

64. Curtsinger, J. W. Late-life fecundity plateaus in Drosophila melanogaster can be explained by variation in reproductive life spans. Exp. Gerontol. 48, 1338–1342 (2013).

65. Potdar, S., Daniel, D. K., Thomas, F. A., Lall, S. & Sheeba, V. Sleep deprivation negatively impacts reproductive output in *Drosophila melanogaster*. J. Exp. Biol. 221, jeb174771 (2018).

66. Ahmed, R. M. et al. Physiological changes in neurodegeneration — mechanistic insights and clinical utility. Nat. Rev. Neurol. (2018). doi:10.1038/nrneurol.2018.23

67. Nilsberth, C. et al. The ‘Arctic’ APP mutation (E693G) causes Alzheimer’s disease by enhanced Abeta protofibril formation. Nat. Neurosci. 4, 887–893 (2001).

68. Fernandez-Funez, P., de Mena, L. & Rincon-Limas, D. E. Modeling the complex pathology of Alzheimer’s disease in Drosophila. Exp. Neurol. 274, 58–71 (2015).

69. Perlis, M. L., Shaw, P. J., Cano, G. & Espie, C. A. Models of Insomnia. 850–866 (2014).

70. Gehrman, P. R., Pfeiffenberger, C. & Byrne, E. M. The role of genes in the insomnia phenotype. Sleep Medicine Clinics 8, 323–331 (2013).

71. Lind, M. J. & Gehrman, P. R. Genetic pathways to insomnia. Brain Sciences 6, (2016).

72. Veatch, O. J., Keenan, B. T., Gehrman, P. R., Malow, B. A. & Pack, A. I. Pleiotropic genetic effects influencing sleep and neurological disorders. The Lancet Neurology 16, 158–170 (2017).

73. Spielman, A., Saskin, P. & Thorpy, M. J. Treatment of Chronic Insomnia by Restriction of Time in Bed. Sleep 10, 45–56 (1987).

74. Perlis, M. L., Corbitt, C. B. & Kloss, J. D. Insomnia research: 3Ps and beyond. Sleep Medicine Reviews 18, 191–193 (2014).

75. Dissel, S. et al. Enhanced sleep reverses memory deficits and underlying pathology in drosophila models of Alzheimer’s disease. Neurobiol. Sleep Circadian Rhythm. 2, 15–26 (2017).

76. Cassidy-Eagle, E. et al. Cognitive Behavioral Treatment for Insomnia in Older Adults with Mild Cognitive Impairment in Independent Living Facilities: A Pilot Study. J. Sleep Disord. Med. Care 1, (2018).

77. Insel, T. R. Next-generation treatments for mental disorders. Sci. Transl. Med. 4, 155ps19 (2012).

78. Hofmann, S. G., Asnaani, A., Vonk, I. J. J., Sawyer, A. T. & Fang, A. The efficacy of cognitive behavioral therapy: a review of meta-analyses. Cogn. Ther. Res. 36, 427–440 (2012).

